# Rats can learn a temporal task in a single session

**DOI:** 10.1101/649954

**Authors:** Marcelo Bussotti Reyes, Diego Henrique de Miranda, Gabriela Chiuffa Tunes, Marcelo Salvador Caetano

## Abstract

Fixed interval, peak interval, and temporal bisection procedures have been used to assess cognitive functions and address questions such as how animals perceive, represent, and reproduce time intervals. They have also been extensively used to test the effects of drugs on behavior, and to describe the neural correlates of interval timing. However, those procedures usually require several weeks of training for behavior to stabilize. Here, we compared three types of training protocols and reported a procedure in which performance by the end of the very first session nearly matches the performance of long-term training. We also discuss this fast-learning protocol in terms of an information-theory approach. This one-day training protocol can be used to investigate temporal learning and may be especially useful to electrophysiological and neuropharmacological studies.

## 1 Introduction

Experimental psychology is the go-to field when it comes to the development and refinement of behavioral tasks, which in turn are used to test hypotheses on a vast range of subjects. Often referred to as schedules of reinforcement (Fester and Skinner 1957), such behavioral protocols have been used in the study of cognitive functions as learning and memory, perception, motivation, and timing, among many others.

There are many behavioral tasks devoted to the study of how animals perceive and reproduce time. Examples are the fixed-interval procedure (Skinner 1938; Roberts and Church 1978), peak procedure (Roberts 1981), temporal bisection (Church and Deluty 1977), gap procedure (Buhusi and Meck 2007), among others. Results obtained from these tasks have been the cornerstone for the expanding field of interval timing, and the source for most of the current knowledge on timing behavior. One common aspect of these procedures is that, in order for humans and non-human animals to achieve relatively stable performance (i.e., asymptotic behavior) in the task, weeks of training are required. Also, those procedures usually deal with judgment or estimation of the duration of a certain stimulus, such as a light or a sound.

There is, however, a subset of protocols that can be trained faster, and for which the duration of the organism’s own response is the temporal interval of interest (such as the time during which a lever was kept pressed, or a nose-poke response was sustained). Skinner (1938, chapter VIII) described this type of schedule as “differentiation of response duration”. Other studies have referred to such a protocol as “response duration differentiation” (RDD, Hudzik et al. 2000), “differential reinforcement of response duration” (DRRD), and “temporal response differentiation” (TRD, McClure, Hardwick, and McMillan 2000). They have been used to describe long-term temporal performance (Mcmillan and Patton 1965; Lejeune and Jasselette 1987; Platt, Kuch, and Bitgood 1973; Kuch 1974; Gulotta and Byrne 2015), the effects of drugs on temporal performance (Bruhwyler, Chleide, and Mercier 1990; McClure, Hardwick, and McMillan 2000; Narayanan and Laubach 2009), in studies of impulsivity (Sanabria and Killeen 2008), brain lesions (Hudzik et al. 2000), and brain encoding (Lebedev, O’Doherty, and Nicolelis 2007). Although used for different purposes, in all studies the protocol took more than five days (sometimes weeks) to be trained. None of them reported significant temporal learning on the first day of training.

Here, we report adjustments in the training protocol which drastically improved the learning performance reported in the literature. We tested three DRRD protocols, assessing the distribution of response duration in the first ten sessions of training. We show that our simplified version of the training protocol not only highly increased the speed of learning but produced significant behavioral changes detected from the very first session. Furthermore, rats achieved asymptotic performance within very few days of training, if not in the very first day. This new protocol is especially suitable for the study of functional and neural mechanisms of temporal learning.

## 2 Materials and Methods

### 2.1 Animals

Eighteen naïve Wistar male rats (purchased from the Federal University of São Paulo, Brazil) were used in the study. They were housed in the university’s vivarium. They were two months old and weighed between 250-350 g at the beginning of the experiment. They were kept in a 12h light/dark cycle (lights on at 7 am). Experimental sessions occurred during the light-on phase. During training, rats were food-restricted and kept between 85 and 90 % of their *ad-libitum* weight. All experimental procedures were approved by the Animal Care and Use Committee at Federal University of ABC (UFABC), and conform to guidelines for the Ethical Treatment of Animals (National Institutes of Health).

### 2.2 Training chambers

Six standard operant chambers (Med-Associates, Inc.), 25 cm wide, 30 cm high, by 30 cm deep, were used during training. On the front wall, each chamber was equipped with a central magazine pellet dispenser, which delivered 45 mg sucrose pellets (Schraibmann LTDA, Brazil) into a food cup, two levers (one at the right and one at the left of the pellet dispenser), and two light stimuli (located above each lever) with diffuse illumination of approximately 200 Lux. The levers were always available throughout the experiment, but only the left lever was used in the training protocol, i.e., presses to the right lever had no consequence. All procedural events were controlled by the MedPC software (Med-Associates, Inc) and recorded with a temporal resolution of 2 ms.

### 2.3 Training protocol

Rats were handled for 2-3 days before training stated. They were randomly assigned to one of three groups: “Timeout”(*n* = 6), “No-Timeout” (*n* = 6), and “Fixed” (*n* = 6). For rats in all groups, the experimental session started with a continuous reinforcement (CRF) schedule, during which every lever press immediately delivered a sucrose pellet. After receiving 100 pellets in less than 45 minutes, each of the three groups of rats was trained in different variations of the Differential Reinforcement of Response Duration (DRRD) protocol (Fig. 1). In general, rats had to sustain a lever-press response for a minimum criterion time, and then release the lever to obtain a sucrose pellet. All rats were trained for ten daily sessions lasting 50 min each, as described below.

**Figure 1:**
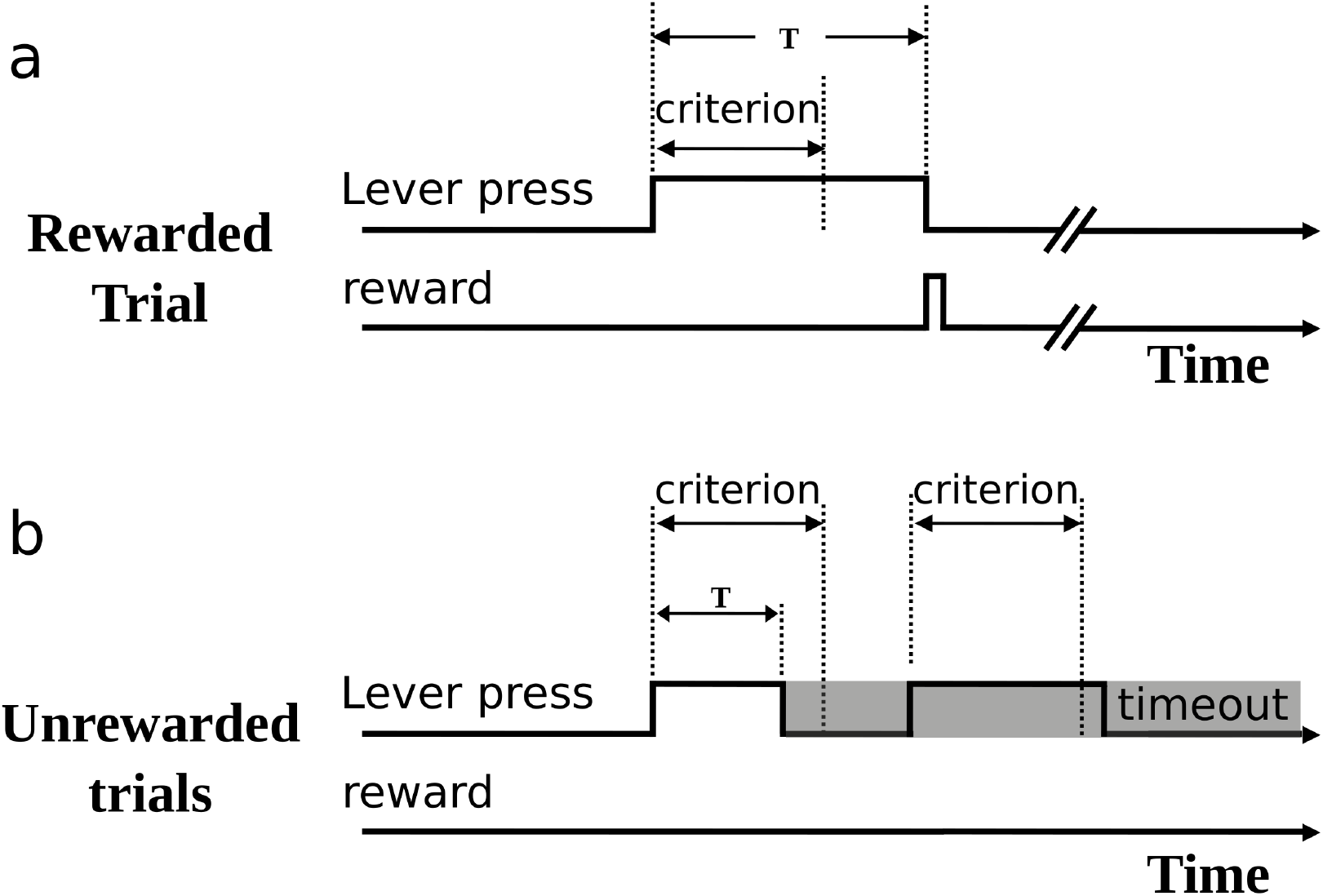
Description of the task. a) Rewarded trials. When the rat held down the lever for an interval longer than the criterion time, a sucrose pellet was delivered upon lever release. b) Unrewarded trials. Lever-press durations shorter than the criterion time were not rewarded and, only for the group Timeout, they were followed by a timeout period (grey rectangle) during which the cue light was turned off. During the timeout period, lever presses restarted the timeout timer and were not rewarded even if their durations were longer than the criterion time.

### 2.4 Group Timeout

The cue light above the left lever was turned on at the beginning of the session to signal the beginning of the trial. The initial criterion in DRRD sessions for this group was 0 ms, i.e., rats had to withhold a lever press to obtain a food pellet (“correct response”). After three consecutive correct responses the criterion increased 100 ms up to 1.2 s, and after six consecutive incorrect responses (shorter than the criterion time), the criterion decreased 100 ms (down to a minimum of 100 ms). The cue light remained on after correct responses but was turned off after incorrect responses for a “timeout” period that varied from 2 to 4 seconds, randomly selected from a uniform distribution. Responses during light off reset the timeout timer were classified as invalid and did not produce any reward regardless of their duration.

### 2.5 Group No-Timeout

For group No-Timeout, training was similar to group Timeout but with three main modifications: The initial criterion was 500 ms. Incorrect responses were not followed by any timeout (i.e., all lever presses were classified as valid, even though only those which were held longer than the criterion time were rewarded); And second, the criterion time never decreased after incorrect responses.

### 2.6 Group Fixed

For group Fixed, the criterion time started at 1.2 s and was kept fixed at this duration throughout training. There were no timeouts (all lever presses were valid) and no increments or decrements in the criterion.

### 2.7 Data analyses

All data were analyzed using Matlab (Mathworks, Inc.) and R (R Core Team 2013) routines developed in our laboratory. We calculated the response distribution by convolving the histogram of the response duration in 100-ms bins with a Gaussian kernel (200 ms width).

## 3 Results

Performance in the DRRD task in the first session developed differently across the three groups. At the beginning of the session, mainly short responses were produced typically around 500 ms. As training in this first session progressed, response duration for some animals shifted from shorter to longer lever presses, increasing their reward rate. Each panel of the Fig. 2 shows a typical performance of each group in the first session. Improvement in performance was analyzed by looking at the distribution of the response duration, i.e., the response counts as a function of time, smothered and normalized. The curves in each plot of Fig. 2 represent the duration distribution in the first 30 trials (dashed line) and the last 30 trials (continuous lines). The animal shown in panel **a** (group Timeout) had little to no improvement in performance, that is, response distributions at the beginning and at the end of the session were comparable. However, the animal from No-Timeout (**b**) and Fixed (**c**) groups showed a clear improvement in performance, displaying many more correct responses at the end of the session, which led to higher reward rates. This improvement is illustrated by a rightward shift of the distribution of responses at the end of the session compared to the beginning.

**Figure 2:**
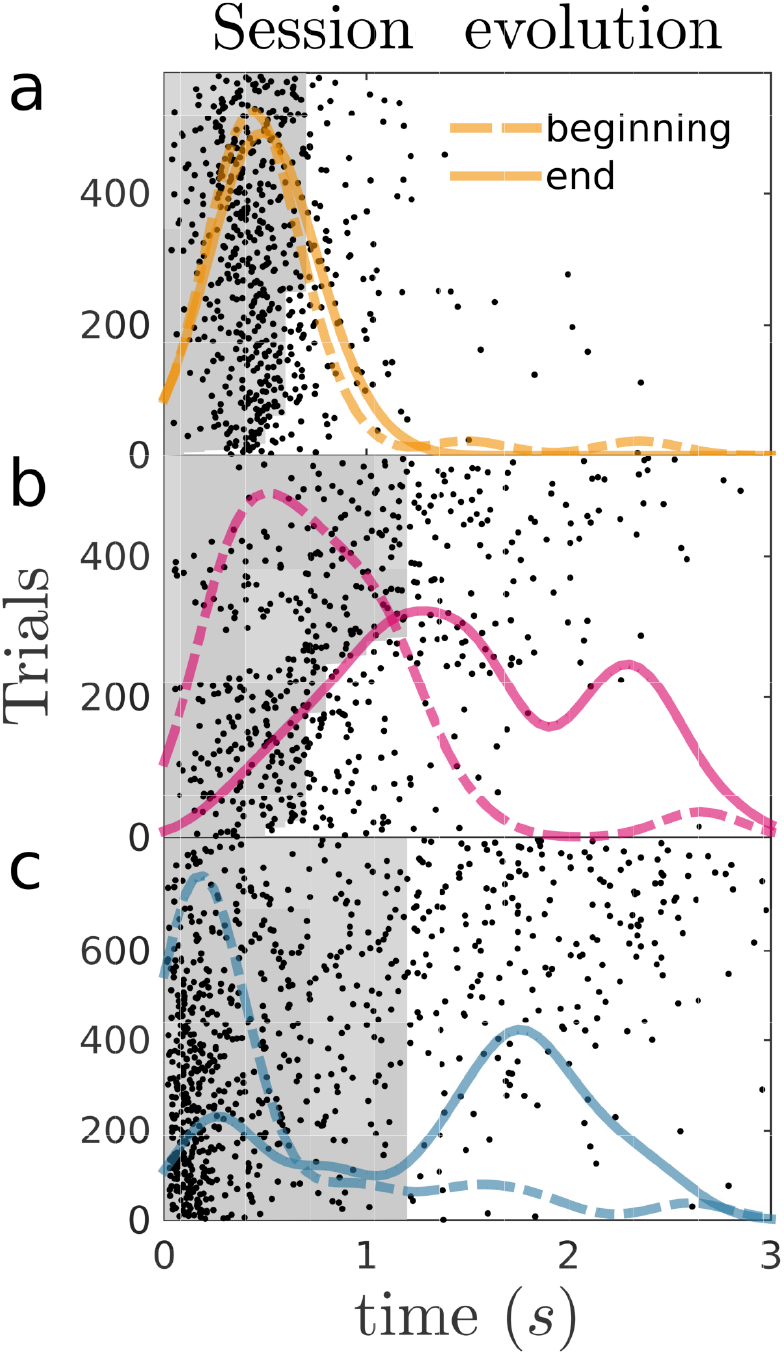
Responses from one rat from each of the three groups: (a) Timeout, (b) No-Timeout, and (c) Fixed. Each black dot represents a nose-poke response duration (x-axis) in each trial of the first session of training (y-axis). The gray shaded area corresponds to the criterion time for each trial—responses within this area were not reinforced. In (a) and (b) the criterion changed over the session (accordingly, the shape of the gray area is irregular), while in (c) the criterion remained constant (Fixed group). The lines show fitted distributions of responses in the first (dashed) and the last (continuous line) 30 trials, normalized and scaled for better visualization.

The differences across groups observed in Fig. 2 came from analyzing a representative rat from each group. To compare the performance of all rats across groups, we pooled 30 response durations at the beginning of the first session, and 30 durations at the end of the session from each animal in each group and compared their distributions (Fig. 3a).

**Figure 3:**
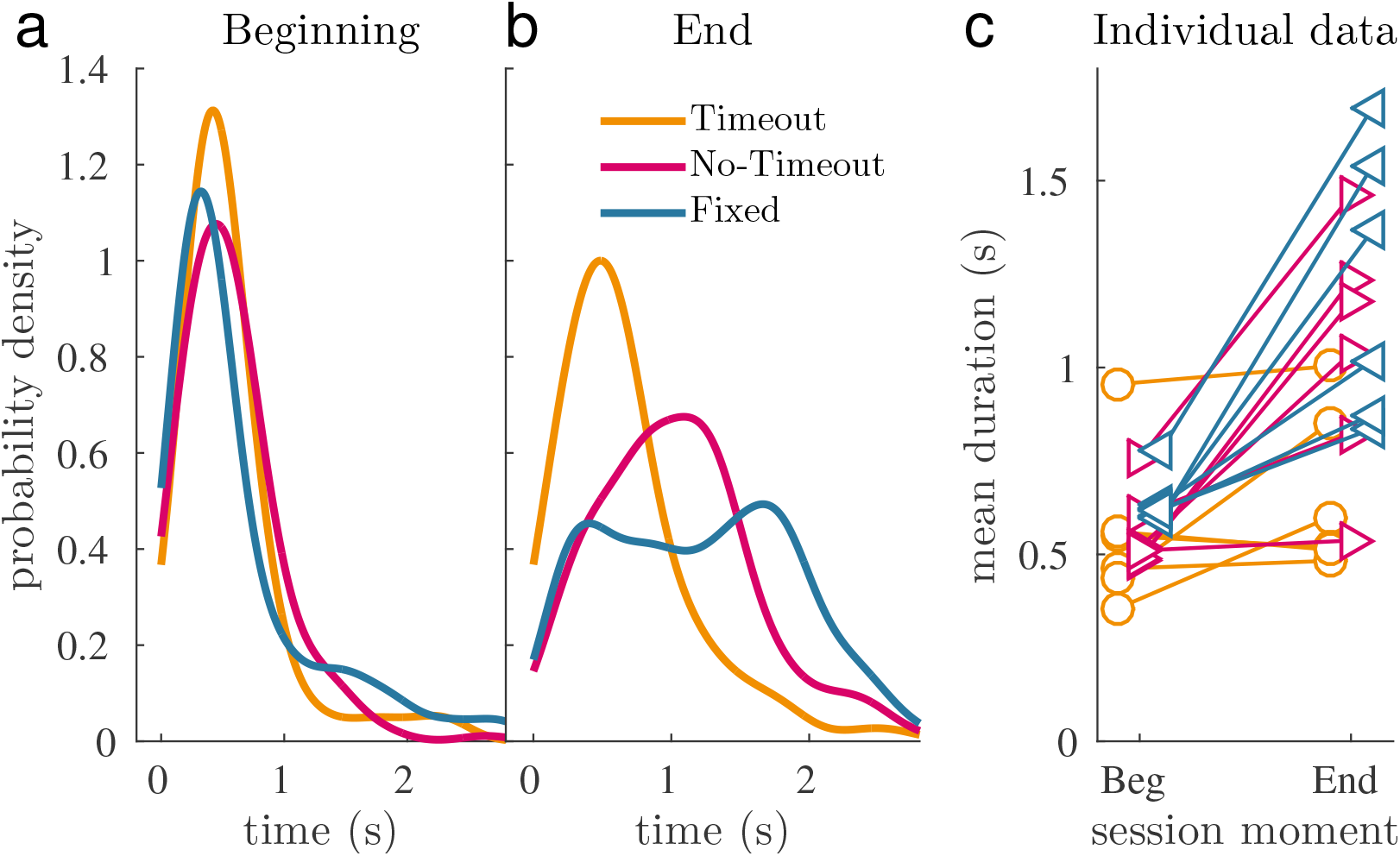
Average performance across groups. a) Probability density functions of response duration at the beginning and at the end of the first session. Distribution of the aggregated data from the first (a) and last (b) 30 trials of all rats in each group (180 total trials per group). The three response distributions peaked at approximately the same time. Right) Distribution of the aggregated data from the last 30 trials of all rats in each group. (180 total trials per group). For the groups No-Timeout and Fixed, the duration distributions shifted to the right. c) Median duration of responses in the first (beg) and last (end) 30 trials from individual animals.

In the beginning, the distributions from the three groups peaked before 500 ms, suggesting that all animals departed from similar baselines. A Kolmogorov-Smirnov comparison showed that distributions from group Timeout and Fixed differed from the groups Timeout and No-Timeout (D=0.16, p=0.022 and D=0.15, p = 0.031, respectively), and the groups Timeout and No-Timeout were similar (D=0.14, p=0.056). Since the Kolmogorov-Smirnov test is very susceptible to the shape of the distributions, and consequently it is not surprising to find significant differences. However, notice that the Kolmogorov distances D were relatively small (around 0.15). By the end of the session (last 30 trials), however, the distributions from the three groups differed as the Kolmogorov statistics yielded higher values (group Timeout versus No-Timeout: D=0.41, p¡0.001; group Timeout versus Fixed: D=0.43, group No-Timeout versus Fixed: D=0.23; p=0.001). These results support that the animals started from similar baselines and the training led them to different duration distributions.

Notice that we compared distributions using the same number of trials per animal and per group. Therefore, any differences observed did not result from a possible bias produced by animals that respond more or less. The curves in Fig. 3a (End) suggest that the distribution of responses for group Fixed had a heavier right tail than for group No-Timeout, possibly producing more long trials in the first session.

To individually describe the difference in behavior produced by the protocols, we compared mean response duration in the first and last 30 trials for each animal between groups (Fig.3c). A Repeated Measures ANOVA analysis showed significant difference across groups (F(2,10)=5.84; p=0.021), effect of beginning vs. end of session (F(1,5)= 31.97; p=0.002), and a significant interaction between group and time in the session (F(2,10)=5.21; p=0.028). This interaction indicates that the groups learned differently in this first session. Hence, we compared each group at the beginning and the end of the first session. In the group Timeout, the mean duration of responses remained constant in the first session (paired t-test, t(5)=1,449,p=0.207), while it varied in the other two groups (group No-Timeout: t(5)=3.9874, p=0.010; group Fixed: t(5)=4.2661, p=0.008). Hence, the learning was different for the group Timeout and the other two.

The difference observed in the first session persisted over consecutive days of training. Throughout the sessions, the mean duration achieved by Group Timeout was clearly lower than those achieved by the other groups. To compare the groups throughout the sessions, for each animal we fitted an exponential curve (*y* = *a − be^−cx^*) to the mean duration of responses of the first and the last 30 trials of the first session, and the mean of the last 30 trials of the consecutive session (2 to 10). Then, to compared the fitted parameters cross the groups through a no parametric test (Krustkal-Wallis). Such comparison showed that there was no evidence that the parameters a and b were different across the groups. However, the parameter c differed between the groups, which means that the exponential had a different rate of change across groups. To assess the difference between groups, we used a Wilcoxon rank sum test for pair-wise comparisons. In agreement with the previous analyses, there was a difference between groups timeout and no timeout (p=0.026) as well as between timeout and fixed (p=0.041), but no difference between the groups fixed and no timeout. Also, only after all ten sessions of training, rats in Group Timeout reached a performance similar to the other two groups. Importantly, by the end of the first session, rats in groups No-Timeout and Fixed had already achieved a performance that was comparable to the asymptotic value of about 1.4 s observed in later sessions, suggesting that in the very first session those animals had already learned the task almost to their best—i.e., learning took place mostly during the first session.

Next, we checked if the behavior after the first session was comparable with the asymptotic behavior (long term performance). We compared the difference of means between the second and last session of each group as a measurement of the improvement in performance. We checked whether such improvement differed from 0 using a t-test. There was no evidence that groups fixed (p=0.23) and no timeout (p=0.91) improved over sessions, while there was a significant improvement for the group timeout (p=0.0095). While these results are inconclusive about whether the groups No-Timeout and Fixed improve after the first session, it clearly shows that the group timeout was still learning.

Finally, we measured temporal learning by assessing how many trials were necessary for rats in each group to produce 100 “long trials” (i.e., response durations longer than 1.2 s). Fig. 5 shows a cumulative record of long trials. Rats in group Timeout on average took more than 1,500 trials to complete 100 long trials, while rats in the other two groups took about 500 trials (Fig.5). There was a significant effect of the number of trials (*f* (2, 15) = 3.79, *p* = 0.046, one-way ANOVA) and HSD posthoc comparisons between groups showed a significant difference between group Timeout and the other two groups (*p* = 0.027 and *p* = 0.035, between group Timeout and group No-Timeout and Fixed, respectively). No differences between groups No-Timeout and Fixed were observed (*p* = 0.89).

## 4 Discussion

We found that one modification in the behavioral protocol—the elimination of a timeout after incorrect responses—increased the learning speed to the point that performance at the end of the first session nearly matched performance after many sessions of training.

To the best of our knowledge, no studies have so far systematically investigated the effect of a timeout in a task of response duration differentiation. The rationale underlying the use of timeout in this task is that it decreases the probability of occurrence of short responses (i.e., those that precede the timeout), in accordance with the law of effect. Moreover, timeouts create a temporal contingency that, in turn, improves the salience of long lever presses—every short response eliminates an opportunity of receiving a reward. However, our results point to the opposite direction: by removing any timeouts and providing no consequences to short responses, the amount of training required for the animals to reach asymptotic performance was considerably reduced. By the end of the very first session, performance of both groups Fixed and No-Timeout was comparable to performance observed after many sessions of training (Fig. 4). On the other hand, rats in Group Timeout took about ten days of training to achieve comparable performance. This larger amount of training is comparable to results in the literature in which the timeout was used (Narayanan and Laubach 2009; Laubach and Pierce 2010).

**Figure 4:**
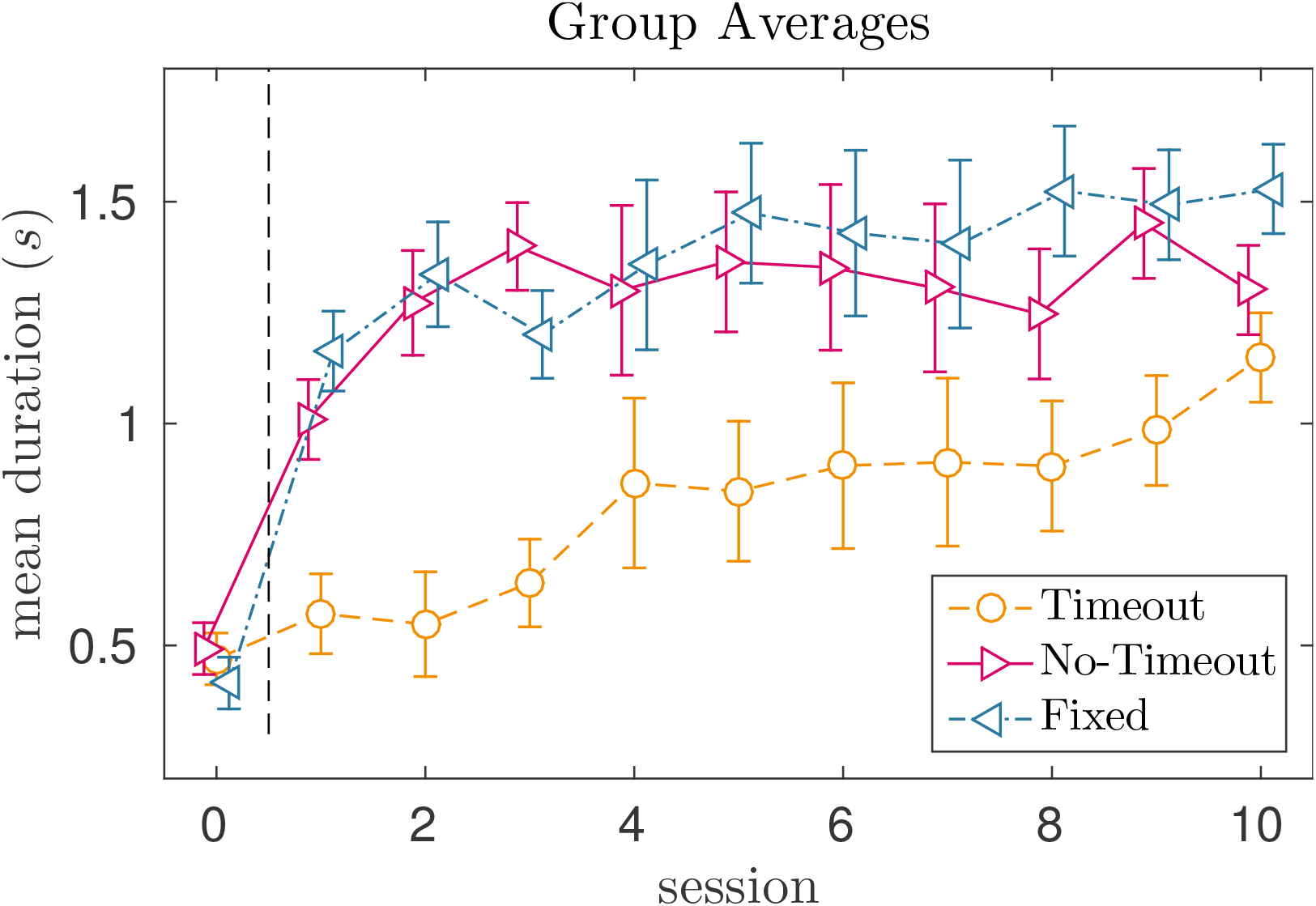
Average response durations in the last 30 trials for each group across sessions. Session 0 shows responses at the beginning of the first session (average durations produced in the first 30 trials). All the other points represent average durations produced in the last 30 trials of all sessions.

**Figure 5:**
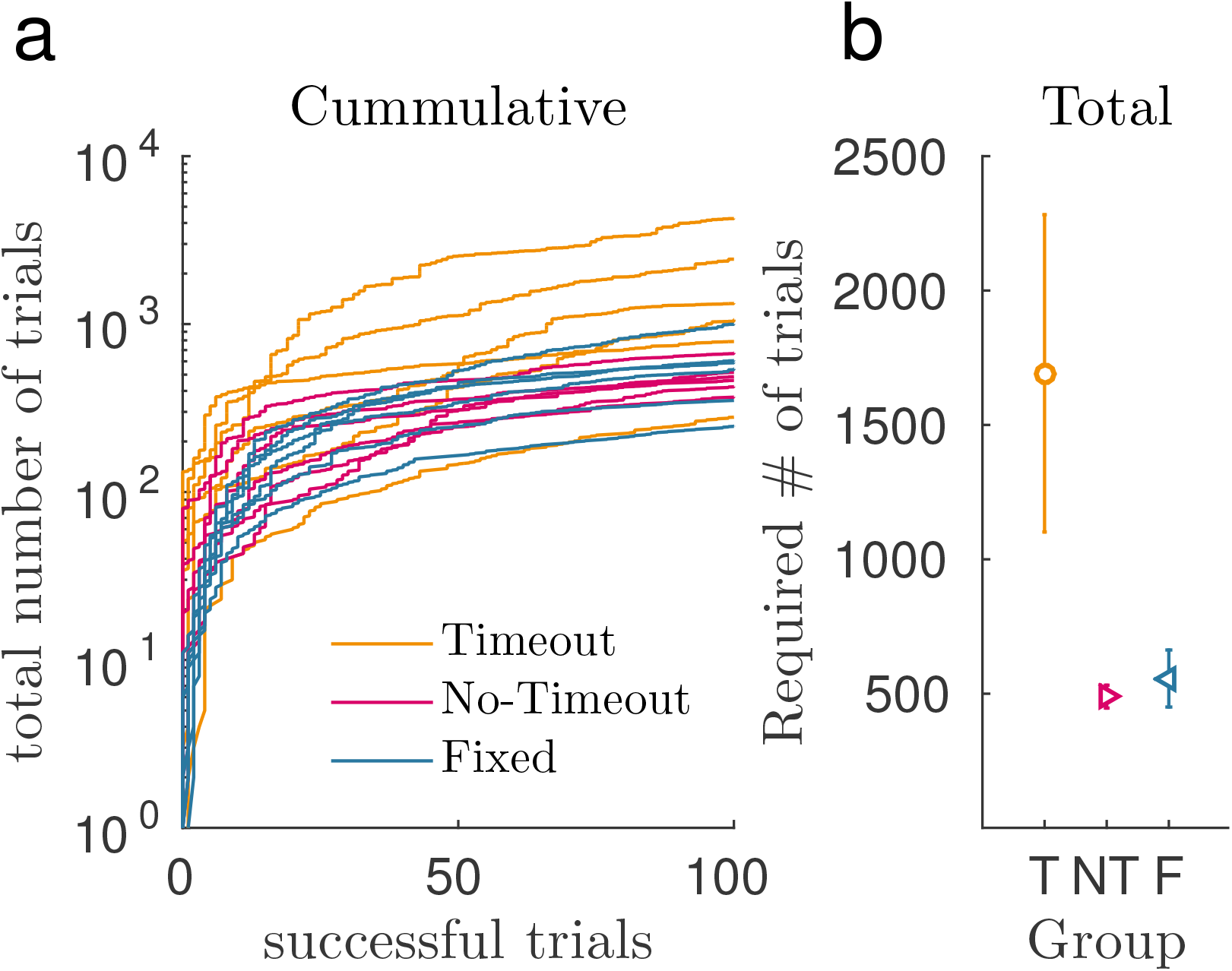
Performance of the individual rats in the first sessions. a) Cumulative of the total number of responses (in a log scale for better visualization) as a function of the number of responses greater than 1.2 *s* (successful responses). b) The total number of responses required to produce 100 successful responses per group. Group labels represent each group: T represents the Timeout, NT the No-Timeout, and F the Fixed group.

Our results are inconclusive as to whether performance between groups No-Timeout and Fixed differed. We did find a significant difference in the number of long trials produced between those groups when we aggregated data from all rats at the end of the first session (i.e, combined the last 30 trials from each rat into one data set per group). The distribution of responses for the Fixed group displayed a longer tail, which suggests more reinforced long trials in this group than in the No-Timeout group. However, when we statistically compared the individual mean response durations between groups, i.e., one measure per rat, that difference did not hold significant. It is possible that individual means do not properly represent the response distribution of each animal. For example, consider two groups of animals, one that produces responses either very short (around 100 ms) or long (around 1.3 s), and another group that responds around 700 *ms* but with a lower variability. These two groups would have the same average response duration, but surely a different rate of rewards. Thus, even though averaging data from different animals may be misleading in some cases, it seems that the difference observed in Fig. 3a is more informative than the statistics from individual data. However, when we compare the number of trials necessary to achieve 100 rewards (Fig. 5), again the groups No-Timeout and Fixed did not differ significantly. Hence, the current experimental paradigm (and perhaps the number of animals trained in each group) does not allow us to tell these two groups apart. Importantly, this does not mean that the two training protocols are equivalent choices for training DRRD schedules: the Fixed protocol has fewer parameters, with no rules for the number of correct responses to increase criterion neither the step size for increments, and therefore it is a simpler and better option for training.

Comparing our results with the literature of response differentiation is difficult because the majority of studies reported training phases between 5 and 10 days long—they were trying to establish a steady performance over sessions, either analyzing the behavior itself (Mcmillan and Patton 1965; Platt, Kuch, and Bitgood 1973; Lejeune and Jasselette 1987) or investigating the effect of drugs on performance (Bruhwyler, Chleide, and Mercier 1990). The focus on the steady performance and its manipulation was probably the reason why previous studies excluded the first sessions from the analyses. Exceptions are the work of Yin and colleagues (Yu, Gupta, and Yin 2010; Fan, Rossi, and Yin 2012) who describe results from the first session and report that some learning occurred. However, they did not analyze intra-session data. We believe we are showing the first study with naive animals within the very first session of training.

The current literature lacks a theoretical framework that predicts which parameters in a protocol produce faster learning. Balsam, Gallistel, and colleagues (Balsam and Gallistel 2009; Ward, Gallistel, and Balsam 2013; Gallistel, Craig, and Shahan 2014) developed a framework that allows us to hint at how fast an organism would learn a task based exclusively on the temporal and probabilistic aspects of the protocol. It states that the amount of training in a given protocol needed for learning to occur is a function of the Shannon information (or a linear combination of entropies) provided by the task. Furthermore, they show how to calculate this information for simple protocols, namely the delay and random protocols in Pavlovian conditioning. They have described how their framework also applies to instrumental learning in a more recent study (Gallistel, Craig, and Shahan 2014). Even though it is still unclear how to calculate the information for a protocol like the DRRD, it seems clear that the fewer contingencies in the task, the smaller the uncertainty about when the food is coming, and hence, the greater the total information. Finally, the higher the information, the faster is learning. Within this framework, the timeout seems to create extra to-be-decoded information, impairing the learning process.

Our results may also inform how to optimize the parameters of the training protocol. For the group Fixed, for example, gauging the size of the criterion is critical: a short criterion would pass unnoticed by the rats while an excessively long criterion—longer than 20 s or 30 s, for example—would be virtually impossible to learn. Therefore, there should be an “ideal” point where learning rate is maximal.

Another relevant aspect of this task is its potential to generate long responses, which are interesting for interval timing studies, for example. Regarding this issue, what is the maximum duration the group No-Timeout (which increments the criterion in steps) can achieve? Gulotta and Byrne (2015) showed that animals were able to keep a lever pressed for about 20 s. And, as argued above, such long responses would not occur in a Fixed protocol. Consequently, there is probably an intermediate protocol, perhaps a combination of the No-Timeout and Fixed, that would generate faster learning of longer responses. Those are still open and exciting questions about this old and yet unexplored aspect of timed behavior.

## Acknowledgements

This study was funded by a grant from the Brazilian National Research Council (CNPq, Grant # 430993/2016-1) to MBR; and by a doctoral fellowship from the São Paulo Research Foundation (FAPESP, Grant # 2015/08132-9) to GCT. MSC is affiliated to Instituto Nacional de Ciência e Tecnologia sobre Comportamento, Cognição e Ensino, supported by CNPq (Grant # 465686/2014-1) and FAPESP (Grant # 2014/50909-8). We would like to thank the members of the Timing and Cognition Laboratory at UFABC (neuro.ufabc.edu.br/timing/) for useful discussions and suggestions on this study.

